# Pollinator specificity among three co-flowering Mediterranean *Aristolochia* species pollinated by Diptera

**DOI:** 10.64898/2026.05.19.726152

**Authors:** Valentin Vrecko, Benoit Lapeyre, Bruno Buatois, Annick Lucas, Raphaelle Aubry, Ryszard Szadziewski, Michael von Tschirnhaus, Aroonrat Kidyoo, Anne-Geneviève Bagnères, Björn Bohman, Doyle McKey, Rumsaïs Blatrix, Magali Proffit

## Abstract

Attracting species-specific pollinators can be favoured by natural selection to avoid reproductive interference between sympatric plant species. However, the ways in which fine differences in floral traits lead to the attraction of species-specific pollinators are diverse and unknown in most pollination interactions. We surveyed pollinators on three *Aristolochia* species that can occur in pairs in sympatry (*A. clematitis*-*A. rotunda* and *A. pistolochia*-*A. rotunda*) and are pollinated by Ceratopogonidae and Chloropidae (Diptera), to investigate the extent to which pollinator assemblages differ among plant species. To assess whether specificity in pollinator assemblages may be mediated by differences in floral odour composition, we characterised the volatile organic compounds (VOCs) emitted by receptive flowers and highlighted those VOCs electrophysiologically detected by pollinators of *A. rotunda* and *A. pistolochia*. Among the most abundant pollinators, female*s of Forcipomyia monilicornis* were specific pollinators of *A. pistolochia* while males and females of two *Dasyhelea* species were specific pollinators of *A. clematitis*. Females of *Forcipomyia aristolochiae* and of *Trachysiphonella ruficeps* were the main pollinators in *A. rotunda* but also pollinated the other two species. The floral odours of *A. rotunda* and *A. pistolochia* differed significantly from each other and elicited specific electrophysiological responses in their respective pollinators. Although several pollinator species visit more than one *Aristolochia* species, those pollinators are preferentially found in one *Aristolochia* species. Selective attraction is likely mediated by specific VOCs.

## Introduction

Although in some cases plant species may benefit from sharing pollinators through the facilitation of pollination (Kemp et al., 2019), in most cases sharing pollinators reduces plant reproductive success through pollen misplacement, nectar robbing, interspecific competition for pollinators and heterospecific pollen deposition (Moreira-Hernández & Muchhala, 2019; Shivanna, 2014). One evolutionary response to avoid reproductive interference in plants is to attract specific pollinators (Moreira-Hernández & Muchhala, 2019). This could be achieved through divergence in floral signals to attract distinct guilds of pollinators in the community (Gervasi & Schiestl, 2017). Even slight changes in floral traits could allow specific attraction of pollinator species different from those of co-flowering plants (Moreira-Hernández & Muchhala, 2019; Muchhala et al., 2014). Deciphering how changes in floral traits lead to specific pollination is particularly relevant among closely related plant species that flower in sympatry and use similar pollination strategies, because selective pressures to avoid reproductive interference may be strong. Unfortunately, how pollinator specificity is modulated by changes in floral phenotype has only been clearly deciphered in very few plant-pollinator interactions, such as the fig-fig wasp obligatory mutualism (Liang et al., 2026; Souto-Vilaros et al., 2018), and sexually deceptive orchid-wasp pollination systems (Johnson & Schiestl, 2016; Schiestl & Schlüter, 2009). To gain a general understanding of how changes in floral traits affect the specificity of pollinator attraction, the range of pollination strategies studied should be expanded.

In recent years, pollination by Diptera has been shown to be far more diverse than previously thought, involving a wide variety of pollinators belonging to over 70 families and a wide range of pollination strategies (Larson et al., 2001; Raguso, 2020). This diversity of interactions is particularly remarkable in certain highly diverse plant taxa that are specialised for pollination by Diptera, such as the genera *Aristolochia* L. (Aristolochiaceae) (Berjano et al., 2009), *Ceropegia* (Apocynaceae) (Heiduk et al., 2015; Ollerton et al., 2017), and *Lepanthes* (Orchidaceae) (Karremans & Díaz-Morales, 2019), among many others. In some of these genera, related plant species growing in sympatry show specificity in their pollinators (often congeneric species of Diptera), which are attracted likely through similar pollination strategies (Chua et al., 2020; Kidyoo et al., 2022; Vogel & Martens, 2000). This makes pollination by flies an excellent system for studying the mechanisms resulting in pollinator specificity among closely related plant species.

The genus *Aristolochia* (Aristolochiaceae) comprises more than 550 accepted plant species, mainly distributed in tropical and Mediterranean regions (Krause et al., 2022; Ohi-Toma et al., 2006). *Aristolochia* species are pollinated exclusively by flies, mostly through floral deception (Knuth, 1909). Although a large diversity of Diptera families was identified as pollinators in the genus, each *Aristolochia* species is generally specialised in flies belonging to only one or two Diptera families (see the review of Berjano et al., 2009; Matallana-Puerto et al., 2025; Oelschlägel et al., 2015; Rupp et al., 2021, 2024). In *Aristolochia*, the morphology of the trap flower varies in association with pollinator morphology (Berjano et al., 2009; Brandalise et al., 2026; Brantjes, 1980; Rulik et al., 2008). Such morphological variation can contribute to pollinator specificity among *Aristolochia* species by favouring pollen deposition on, and floral access by, pollinators of appropriate size (Berjano et al., 2009; Brandalise et al., 2026; Bröderbauer et al., 2013). However, morphological filtering has been shown in other trap flowers to be insufficient to ensure pollinator specificity when potential pollinators are morphologically very similar (Kidyoo et al., 2022). Instead, specific pollinator attraction seems to provide a more effective mechanism for maintaining pollinator specificity in deceptive pollination systems involving flies (Kidyoo et al., 2022).

In *Aristolochia*, as in deceptive fly pollination generally (Hayashi et al., 2021; Heiduk et al., 2015; Mochizuki, 2025), olfactory attraction by volatile organic compounds (VOCs) plays an important role in pollinator attraction (Hall & Brown, 1993; Oelschlägel et al., 2015; Rupp et al., 2024). Floral VOCs have been characterised in several *Aristolochia* species: *A. bianorii* Sennen & Pau (Alpuente et al., 2023), *A. baetica* L. (Rupp et al., 2024), *A. cymbifera* Mart. (Johnson & Jürgens, 2010), *A. fimbriata* Cham. (Qin et al., 2021), *A. gigantea* Mart. & Zucc. (Martin et al., 2017), *A. microstoma* Boiss. & Spruner (Rupp et al., 2021), *A. rotunda* L. (Oelschlägel et al., 2015), and *A. ringens* Vahl (Stashenko et al., 2009). But VOCs mediating pollinator attraction have so far been identified in only *A. rotunda* and *A. baetica*, both native to the Mediterranean region. Very different VOCs were characterised among *Aristolochia* species depending on the deception strategy deployed by flowers and on the mimicked models with which they are associated (Johnson & Jürgens, 2010; Oelschlägel et al., 2015; Rupp et al., 2024; Vrecko et al., 2025).

In the Mediterranean region, the sympatric *A. baetica* and *A. paucinervis* Pomel were pollinated by different set of Drosophilidae and Phoridae (Berjano et al., 2009). However, the pollinator species were not always identified to the species level, and the floral mechanisms driving the preferential attraction of distinct pollinators by the two plant species were not investigated. To examine pollinator specificity among *Aristolochia* species in greater detail, we focused on three Mediterranean species of *Aristolochia* that co-occur in the same habitats in southern France with overlapping flowering periods (synchronopatry). These species were *A. rotunda* L.*, A. pistolochia* L., and *A. clematitis* L. *Aristolochia rotunda* is known to attract a kleptoparasitic fly, *Trachysiphonella ruficeps* Macquart (Chloropidae), as well as Ceratopogonidae, by mimicking the odour of injured bugs (Miridae) (Oelschlägel et al., 2015). The flowers of *A. clematitis* and *A. pistolochia* are known to respectively attract Ceratopogonidae and various flies belonging to the Nematocera (Daumann, 1971; Pérez Chiscano, 2011). However, the identity of their effective pollinators is still unknown. If these different sympatric *Aristolochia* species are pollinated by the same families of flies and employ similar pollination strategies, there may be a risk of reproductive interference between them (via heterospecific pollen transfer and/or other mechanisms). These three *Aristolochia* species constitute, then, an appropriate biological model to study pollinator specificity among congeneric plant species and the underlying mechanisms. The aim of this work is to determine whether pollinator specificity occurs among synchronopatric *Aristolochia* species involved in pollination by Diptera, and if so, to determine precisely how differences in floral odours and pollinator-detected VOCs are likely to mediate specific (or selective) pollinator attraction. We thus address the following questions: (1) Does fruit set in *A. clematitis*, *A. pistolochia* and *A. rotunda* depend exclusively on pollinators visiting the flowers? (2) What is the composition of the pollinator communities of these *Aristolochia* species and does each species have specific pollinators? (3) Does pollinator specificity still occur in the context of synchronopatry? (4) Are there differences in the emitted floral VOCs among these species? (5) Do pollinators preferentially detect the floral VOCs emitted by the *Aristolochia* species they pollinate, thereby explaining their specific attraction?

## Material and Methods

### Study site and species

The study was conducted between 2020 and 2025 in 35 collection sites within a radius of 35 km from Montpellier, France (N43.62364, E3.86198), located in the Mediterranean region (Figure S1). The collection sites reflect the patchy distribution of *Aristolochia* individuals in the studied area. The genus *Aristolochia* possesses protogynous flowers, i.e., female receptivity occurs before pollen release. Flowers of *Aristolochia* are characterised by a pronounced syntepaly, forming a chamber, called the utricle, with a perianth tube surmounted by the limb (Oelschlägel et al., 2009). Epicuticular waxes on the limb and downward-pointing hairs along the perianth tube lead pollinators to fall into the flower and be trapped inside (Knuth, 1909). At anther dehiscence, these conical hairs wilt, allowing the pollinator to leave the flower, dusted with pollen. In the study region, the three studied *Aristolochia* species have a flowering period from the end of March to the end of June. In addition, *A. clematitis* and *A. pistolochia* have a secondary period from the end of September to mid-November. In the wettest habitats, flowering of *A. clematitis* may persist during the summer (June to August).

### Reproductive strategy

To determine whether fruit set depends on insect pollination, we quantified fruit set under pollinator exclusion (i.e., through autonomous self-fertilisation or apomixis) on potted plants in the experimental field of CEFE (Centre d’Ecologie Fonctionnelle et Evolutive, Montpellier, France, N43.63861, E3.86361). Flower buds of *A. rotunda* (N = 28) and *A. pistolochia* (N = 15) were bagged with white nylon tissue (mesh ≃ 0.2 mm), allowing light and air to circulate but preventing the entrance of pollinators. For *A. clematitis*, freshly opened flowers (N = 25) were sealed with a small piece of cotton inserted into the floral tube in order to prevent entrance of pollinators. This was preferred over bagging because of the verticillate structure of the axial inflorescence and the short, fragile pedicel of the flower in this species. A maximum of four freshly opened flowers were sealed on the same individual plant, always on different stems. For the three studied *Aristolochia* species, plant individuals have commonly from one to four stems. As a proxy for cross-pollination, fruit set was quantified under hand pollination with allopollen. We did not use natural pollination as a control because the frequency of pollinator occurrence is variable and unpredictable. Hand pollination consisted in making a slit on the side of the utricle with a scalpel and inserting and depositing on the stigmas three pairs of anthers excised from male-stage flowers from different individual plants (Blatrix et al., 2024). Hand pollination was performed on 20 flowers of *A. rotunda,* 12 of *A. pistolochia* and 27 of *A. clematitis*. No more than two flowers were hand-pollinated on the same individual plant. Floral bud bagging and hand pollination were not performed on the same individual plants. To avoid effects of potential reallocation of resources according to fertilisation status (Knight et al., 2006), non-experimental flowers and fruits initiated from them were removed. After five weeks, fruiting success was quantified by noting the presence or absence of fruit. The effect of the treatment (autonomous self-fertilisation or hand pollination) on fruit set (absence or presence of fruit) was tested separately for each *Aristolochia* species by constructing binomial Generalized Linear Models (GLMs; *glm* function in R). Because complete separation occurred in some species–treatment combinations due to the absence of fruit set in one treatment, bias-reduced logistic regression was used to obtain finite parameter estimates, using the *brglm2* package (Kosmidis & Firfth, 2021).

### Identity of floral visitors

The presence of floral visitors within the trap flowers was detected by shining light from a powerful LED (in a smartphone or compact camera) against the utricle, revealing the presence or absence of insects through transparency. Flowers occupied were collected and dissected in the lab under a stereomicroscope, and the floral visitors were preserved in ethanol. The stage of the flower (female or male) and the sex of each floral visitor were recorded. Floral visitors were categorised as “pollinators” if they were carrying pollen in a female-stage flower and pollen was also present on the stigmas, as “pollen carriers” if they were carrying pollen in a male-stage flower or in a female-stage flower without pollen on its stigmas, and as “undetermined” in all other cases. A species of floral visitor was considered as a pollinating species when at least one individual was categorised as “pollinator”. A pollinating species was considered a specific pollinator for an *Aristolochia* species if pollinators (individuals carrying pollen in a female-stage flower and pollen was also present on the stigmas) were found only in this *Aristolochia* species. For individuals carrying pollen in female-stage flowers, we counted the number of pollen grains on the body as well as of those deposited on the stigmas of the flower (number was estimated to the nearest 50 if the total was higher than 100) and we summed up the two. The pollen deposited on the stigma was divided equitably among the different “pollinators” if more than one pollinator species was found in a given flower.

For 143 sampling sessions (we consider one session as the survey of a given site for a given date and for a given species of *Aristolochia*) out of 179 (79.9%), we counted the number of flowers inspected, so that the frequency of flowers occupied by pollinators could be computed. Individuals of the families Chloropidae and Ceratopogonidae (the two families that included pollinators) were identified at the species level or classified into morphospecies using morphological characters under the stereomicroscope for Chloropidae and under the microscope on slide-mounted specimens for Ceratopogonidae.

### Sex and reproductive status of the pollinators

If the three *Aristolochia* species employ a similar strategy to attract Diptera, they should attract pollinators with similar characteristics of sex and reproductive status. To obtain the proportion of gravid females of *Forcipomyia* and *Dasyhelea* (Ceratopogonidae), we determined the reproductive status of 21 females (*Forcipomyia*) found in the flowers of each of *A. rotunda* and *A. pistolochia* and of 50 females (*Dasyhelea*) found in the flowers of *A. clematitis.* Presence of chorionated eggs or large oocytes was determined by dissecting the abdomen of females under the stereomicroscope. Presence of eggs was uncertain in some females coming from *A. clematitis*, so the proportion of gravid females was calculated twice, considering eggs as absent or present, respectively, in these ‘uncertain’ females.

To verify that the females of Ceratopogonidae attracted by the flowers of *A. rotunda* had already mated with males (in line with the hypothesis of a pollination strategy based on insect-host mimicry (see Vrecko et al., 2025), the spermathecae of six females of *Forcipomyia aristolochiae* and of one female of another (unidentified) species of *Forcipomyia*, all found in the flowers of *A. rotunda,* were dissected, slide-mounted and observed under the microscope.

Finally, to determine if females were more attracted to flowers than males were, we tested, using binomial tests, the difference between a balanced sex ratio (1 : 1, the typical adult sex-ratio encountered in Ceratopogonidae (Kirk-Spriggs & Sinclair, 2017)) and the number of males and females observed visiting flowers, for each pollinator species of each *Aristolochia* species.

### Factors of variation in pollinator communities

Three parameters were identified as potential factors explaining the composition of pollinator communities: the identity of the *Aristolochia* species, the geographic location and the sampling season. Only the fly species classified as pollinators were included in the community data to focus on the range of floral visitors that effectively exert selective pressure on the floral traits of the plants. First, we tested for a spatial effect on pollinator communities with a Procrustes analysis (Peres-Neto & Jackson, 2001) for each *Aristolochia* species using two distance matrices: one based on the Bray-Curtis dissimilarity index (Bray & Curtis, 1957) of pollinator communities between flowers, and the other based on geographical dissimilarity using the Haversine distance index calculated using GPS coordinates of the sampled populations. Second, we tested the effects of the three parameters on pollinator communities with a Generalised linear model for Multivariate Abundance Data (Multivariate GLM) (function manyglm, *mvabund)* (Wang et al., 2012) based on a negative binomial distribution. Sampling data were pooled by “sampling session”, i.e., combination of *date* x *location* x *plant species,* and a pollinator community matrix was constructed. The plant species, the month of collection, and the collection site (as a categorical factor) were defined as fixed effects.

### Comparisons of floral visitors in synchronopatry

For a more in-depth test of specificity in pollinator attraction among the three *Aristolochia* species studied, we considered only collection sites where at least two species of *Aristolochia* were found in synchronopatry, because pollinators in these sites are in a situation of choice between flowers of different species, excluding confounding factors such as variation in spatial distribution and phenology among species of pollinators. We focused on two synchronopatric situations: *A. rotunda* + *A. pistolochia* (eight sites) and *A. rotunda* + *A. clematitis* (five sites) (*A. pistolochia* and *A. clematitis* were not found in sympatry). Sampling sites were selected when the abundance of flowers of the two *Aristolochia* species was high and visually relatively balanced, thereby limiting the potential influence of the relative abundance of the two plant species on pollinator choice. For each session in these sites, we sampled pollinators in the two *Aristolochia* species simultaneously and recorded the total number of flowers investigated in each species. To avoid overcounting insects due to aggregation in a flower, we used the number of flowers occupied by at least one individual of a pollinator species as a proxy for the species’ abundance. For each species of pollinator and each synchronopatric situation, we pooled data from all sessions in which this species of pollinator was found, and computed χ^2^ tests of independence using the number of flowers occupied and the number of flowers investigated of each *Aristolochia* species.

### Floral VOCs collection and analysis

To investigate whether differences in floral odour composition could modulate specific pollinator attraction, we focused on *A. rotunda* and *A. pistolochia*, two species pollinated by non-gravid females of *Forcipomyia* (Ceratopogonidae) that may rely on similar deceptive pollination strategies. Floral VOCs were sampled in the field using dynamic headspace sampling (see Supporting Method 1.a). Volatiles were captured by trapping on ChromatoProbe® quartz microvials (Varian Inc., length: 15 mm; inner diameter: 2 mm), filled with a mixture of 1.5 mg Tenax-TA and 1.5 mg Carbotrap (60–80 and 20–40 mesh respectively, Sigma-Aldrich, Munich, Germany). Flowers (with stems and leaves) were bagged directly on the living plant. Sixteen and seven individuals, each bearing three to nine receptive flowers, were sampled for *A. pistolochia* (n = 2 at one site and n = 14 at a second site) and *A. rotunda* (n = 7, all at the same site), respectively. Sampling using non-flowering stems was also performed for each plant species (control samples). Bagged flowers were left for 30 min without extraction to ensure concentration of the VOCs inside of the bag. Then, air was drawn out of the bag and through the adsorbent trap for 5 min at a flow rate of 250 ml min^−1^. This sequence of accumulation-extraction was repeated twice for each sample (see Supporting Method 1.a for the details).

The VOC samples were analysed by gas chromatography-mass spectrometry using a semi-standard non-polar capillary column (Optima 5-MS) (see Supporting Method 2). Annotation of VOCs was performed by comparing mass spectra and retention indices with those from a database (NIST 2007 MS library, Wiley 9th edition and Adams, 2007). Confirmation of identity by comparison of retention time and mass spectra with those of synthetic references in our possession was performed for all alkanes, some terpenes and for three electrophysiologically active alkenes that we synthesised: (*Z*)-Pentadec-7-ene, (*Z*)-Pentadec-5-ene and (*Z*)-Heptadec-8-ene. VOC integration was done on the Total Ion Current Chromatogram (TICC) using the MZMine software (version 3.3) (see Supporting Method 2 for details on the integration parameters). Peak areas of the controls (stems and leaves) were subtracted from those of the floral samples (stems, leaves and flowers). This may result in biases in the relative proportions of attractive VOCs of those that are shared between leaves and flowers. As we expect floral VOCs that are important for the attraction of pollinators to be detected in most of the individuals, compounds were retained for analysis only if they were detected in at least 50% of the samples of any one species, independently of the sampling site. The threshold to consider a compound as present was fixed at 0.1% of the total area of the VOCs detected from a sample. Compounds present in quantities under this threshold were not taken into account in the multivariate analyses and were categorised as “trace amount”. The double-bond positions of the different alkenes were identified by analysing the diagnostic fragmentation ions obtained after dimethyl disulfide (DMDS) derivatisation of hexane floral extracts from *A. pistolochia* and *A. rotunda* (Carlson et al., 1989). Details of the preparation of the floral extracts and of the derivatisation protocol and analyses are provided in Supporting Methods 1.b and 3.

To determine if there were significant differences in the floral bouquets of *A. rotunda* and *A. pistolochia*, relative proportions of the compounds were centered log-ratio transformed (clr). Then, the effect of the plant species was tested statistically using a partial least square-discriminant analysis (PLS-DA) followed by a permutation test based on cross-model validation, using the package *RVAideMemoire* (Hervé, 2025).

### Pollinator-antennal olfactory detection

To determine which VOCs are electrophysiologically detected by antennae of *Forcipomyia aristolochiae* and *F. monilicornis* (the main pollinators of *A. rotunda* and *A. pistolochia*, respectively, according to our results), Gas Chromatography - Electroantennographic Detection (GC-EAD) analyses were conducted using natural floral odour extracts prepared following the method of Proffit et al. (2020), using an adsorption-desorption headspace technique (see Supporting Method 1.b for preparation protocol). Analyses were conducted on pollinators on the day of their collection in flowers (see Supporting Method 4). In total, six females of *F. monilicornis* were tested with the floral odours of *A. pistolochia*, and five females of *F. aristolochiae* were tested with the floral odours of *A. rotunda*. In order to investigate which VOCs in the blend of *A. rotunda* could be detected by the main pollinator of *A. pistolochia*, five assays on females of *F. monilicornis* were done using the floral odour of *A. rotunda*. A compound was considered as electrophysiologically active when it elicited a measurable depolarisation in at least half of the individuals tested.

### Field bioassays

To test whether attraction of floral visitors was based on chemical signals, olfactory attraction was tested in the field using McPhail traps. A McPhail trap consists of a transparent plastic container with a central opening at the bottom. Flying insects entering the container are attracted to the top by skylight, and thus, are trapped. A small glass vial containing a piece of paper receiving the odour attractant to test is placed in the trap (Steyskal, 1977). The traps were set for 30-40 minutes at various locations around Montpellier throughout the flowering season. In total, 127 traps were set, in which we deposited between 10 and 40 microliters of hexyl butanoate at different concentrations. We chose to test olfactory attraction by hexyl butanoate because this compound was considered to be the main floral attractant of *A. rotunda* (Oelschlägel et al., 2015), and the field bioassays were conducted before our own analyses of the floral VOCs. Hexyl butanoate was diluted in acetone to speed up the evaporation of the solvent in the traps. In addition, 60 McPhail traps were set without an odour attractant as a control, and 13 traps were set with only acetone to verify that neither the solvent nor the visual traits triggered a behavioural attraction.

## Results

### Reproductive strategy

No fruits were produced under the pollinator-exclusion treatment for *A. clematitis* (N = 25 flowers) or for *A. rotunda* (N = 28 flowers), while only one fruit was produced for *A. pistolochia* (15 flowers, 6.67%). Fruit set was significantly higher under the hand-pollination treatment for the three *Aristolochia* species, with eight fruits produced out of 12 flowers (67%) for *A. pistolochia* (β = 2.90 ± 1.07, z = 2.7, p = 0.007), eight fruits produced out of 20 flowers (40%) for *A. rotunda* (β = 3.66 ± 1.52, z = 2.4, p = 0.016) and 10 fruits produced out of 27 flowers (37%) for *A. clematitis* (β = 3.42 ± 1.51, z = 2.266, p = 0.023).

### Identity of floral visitors

We collected 685 floral visitors from 465 occupied flowers of *A. clematitis*, 668 from 257 occupied flowers of *A. pistolochia* and 1085 from 785 occupied flowers of *A. rotunda*. Most floral visitors (*A. clematitis*: 90.4%; *A. pistolochia*: 94.9%; *A. rotunda*: 88.11%) and all the effective pollinators were of the order Diptera. Floral visitors belonged to the dipteran families Ceratopogonidae (32.8%; 83.8%; 48.1%, at *A. clematitis*, *A. pistolochia*, and *A. rotunda*, respectively), Chloropidae (9.4%; 3.7%; 22.5%), Cecidomyiidae (26.4%; 0.45%; 1.8%) and a few other less represented ones (Figure S2). Pollinating species were restricted to families Ceratopogonidae and Chloropidae. In *A. rotunda*, *Forcipomyia aristolochiae* (Ceratopogonidae) accounted for 68.7% of the pollinators sampled and 79.2% of the pollen transferred. *Trachysiphonella ruficeps* (Chloropidae), accounted for only 12% of the pollinators sampled and 17.8% of the pollen transferred. For *A. pistolochia, F. monilicornis* (Ceratopogonidae) accounted for 74.2% of the pollinators sampled and 65.9% of the pollen transferred, followed by *F. aristolochiae* accounting for 22.5% of the pollinators sampled and 33.6% of the pollen transferred. In *A. clematitis*, *Dasyhelea turficola* and *D. punctiventris* (Ceratopogonidae) accounted respectively for 37.5% and 25% of the pollinators sampled and 43% and 21.2% of the pollen transferred. The other occasional pollinators were three species of Ceratopogonidae and six species of Chloropidae for *A. rotunda*, one species of Chloropidae for *A. pistolochia*, and three species of Ceratopogonidae for *A. clematitis* (Table 1). Among the pollinating species (listed in Table 1), the proportion of specimens detected as pollinators (i.e., found carrying pollen in female-stage flowers with pollen on the stigmas) was much lower in *A. clematitis* (6 %) and *A. pistolochia* (5 %) than in *A. rotunda* (21 %). *Forcipomyia aristolochiae*, the main pollinator of *A. rotunda*, showed a relatively high proportion of specimens detected as pollinators (26 %) in this plant species, compared to the main pollinators of *A. pistolochia* (*F. monilicornis*, 10 % of the specimens detected as pollinators) and *A. clematitis* (*Dasyhelea* spp., 13 % of the specimens detected as pollinators).

**Table 1:**
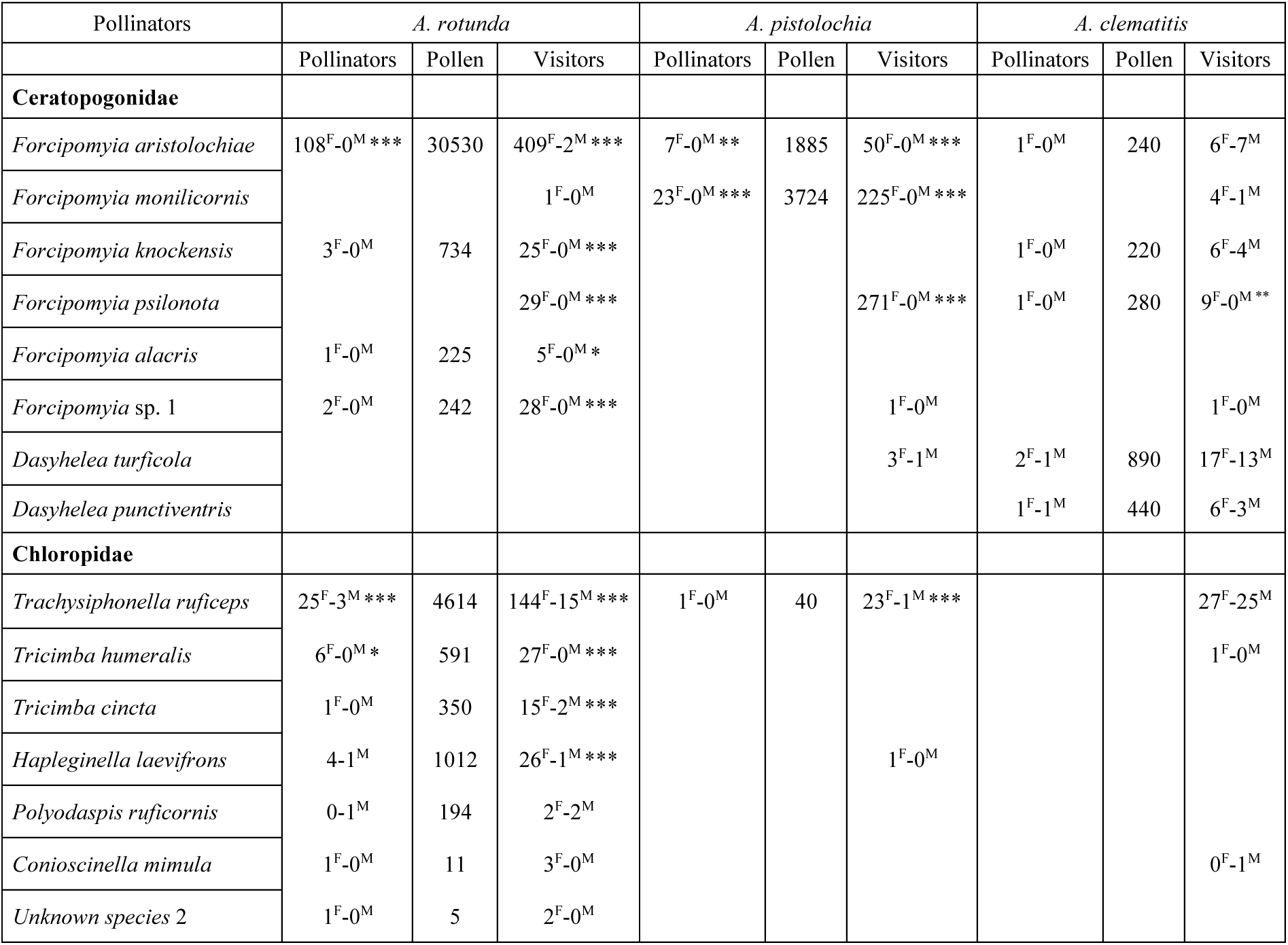
Characteristics of the pollinators of *Aristolochia clematitis*, *A. pistolochia* and *A. rotunda*. Pollinators: the number of floral visitors classified as pollinators (i.e., found carrying pollen in female-stage flowers with pollen on the stigmas). Pollen: total number of pollen grains carried by all the individuals classified as pollinators and those deposited on the stigmas. Visitors: total number of individuals of the pollinator species found in the flowers. For the binomial tests, the observed numbers of males (M) and females (F) of each pollinator species were tested against an expected 1:1 sex ratio in each *Aristolochia* species (*** P < 0.001, ** P < 0.01, * P < 0.05).

| Pollinators | *A. rotunda* |  |  | *A. pistolochia* |  |  | *A. clematitis* |  |  |
|---|---|---|---|---|---|---|---|---|---|
|  | Pollinators | Pollen | Visitors | Pollinators | Pollen | Visitors | Pollinators | Pollen | Visitors |
| **Ceratopogonidae** |  |  |  |  |  |  |  |  |  |
| *Forcipomyia aristolochiae* | 108<sup>F</sup>-0<sup>M</sup> *** | 30530 | 409<sup>F</sup>-2<sup>M</sup> *** | 7<sup>F</sup>-0<sup>M</sup> ** | 1885 | 50<sup>F</sup>-0<sup>M</sup> *** | 1<sup>F</sup>-0<sup>M</sup> | 240 | 6<sup>F</sup>-7<sup>M</sup> |
| *Forcipomyia monilicornis* |  |  | 1<sup>F</sup>-0<sup>M</sup> | 23<sup>F</sup>-0<sup>M</sup> *** | 3724 | 225<sup>F</sup>-0<sup>M</sup> *** |  |  | 4<sup>F</sup>-1<sup>M</sup> |
| *Forcipomyia knockensis* | 3<sup>F</sup>-0<sup>M</sup> | 734 | 25<sup>F</sup>-0<sup>M</sup> *** |  |  |  | 1<sup>F</sup>-0<sup>M</sup> | 220 | 6<sup>F</sup>-4<sup>M</sup> |
| *Forcipomyia psilonota* |  |  | 29<sup>F</sup>-0<sup>M</sup> *** |  |  | 271<sup>F</sup>-0<sup>M</sup> *** | 1<sup>F</sup>-0<sup>M</sup> | 280 | 9<sup>F</sup>-0<sup>M</sup> <sup>**</sup> |
| *Forcipomyia alacris* | 1<sup>F</sup>-0<sup>M</sup> | 225 | 5<sup>F</sup>-0<sup>M</sup> * |  |  |  |  |  |  |
| *Forcipomyia* sp. 1 | 2<sup>F</sup>-0<sup>M</sup> | 242 | 28<sup>F</sup>-0<sup>M</sup> *** |  |  | 1<sup>F</sup>-0<sup>M</sup> |  |  | 1<sup>F</sup>-0<sup>M</sup> |
| *Dasyhelea turficola* |  |  |  |  |  | 3<sup>F</sup>-1<sup>M</sup> | 2<sup>F</sup>-1<sup>M</sup> | 890 | 17<sup>F</sup>-13<sup>M</sup> |
| *Dasyhelea punctiventris* |  |  |  |  |  |  | 1<sup>F</sup>-1<sup>M</sup> | 440 | 6<sup>F</sup>-3<sup>M</sup> |
| **Chloropidae** |  |  |  |  |  |  |  |  |  |
| *Trachysiphonella ruficeps* | 25<sup>F</sup>-3<sup>M</sup> *** | 4614 | 144<sup>F</sup>-15<sup>M</sup> *** | 1<sup>F</sup>-0<sup>M</sup> | 40 | 23<sup>F</sup>-1<sup>M</sup> *** |  |  | 27<sup>F</sup>-25<sup>M</sup> |
| *Tricimba humeralis* | 6<sup>F</sup>-0<sup>M</sup> * | 591 | 27<sup>F</sup>-0<sup>M</sup> *** |  |  |  |  |  | 1<sup>F</sup>-0<sup>M</sup> |
| *Tricimba cincta* | 1<sup>F</sup>-0<sup>M</sup> | 350 | 15<sup>F</sup>-2<sup>M</sup> *** |  |  |  |  |  |  |
| *Hapleginella laevifrons* | 4-1<sup>M</sup> | 1012 | 26<sup>F</sup>-1<sup>M</sup> *** |  |  | 1<sup>F</sup>-0<sup>M</sup> |  |  |  |
| *Polyodaspis ruficornis* | 0-1<sup>M</sup> | 194 | 2<sup>F</sup>-2<sup>M</sup> |  |  |  |  |  |  |
| *Conioscinella mimula* | 1<sup>F</sup>-0<sup>M</sup> | 11 | 3<sup>F</sup>-0<sup>M</sup> |  |  |  |  |  | 0<sup>F</sup>-1<sup>M</sup> |
| *Unknown species* 2 | 1<sup>F</sup>-0<sup>M</sup> | 5 | 2<sup>F</sup>-0<sup>M</sup> |  |  |  |  |  |  |

### Sex and reproductive status of the pollinators

Pollinators of *A. pistolochia* and *A. rotunda* were almost exclusively females (Table 1). In contrast, for *A. clematitis*, the sex ratio was balanced for most of the pollinator species. No gravid females of *Forcipomyia* were found among the 21 dissected females collected in the flowers of *A. rotunda* or among the 21 collected in the flowers of *A. pistolochia*. For *A. clematitis*, the presence of eggs could not be confirmed in 12 (24%) of the 50 *Dasyhelea* individuals examined. Thus, the proportion of gravid females may range from 0 to 12 (0–24%) out of 50 individuals. Sperms were found in the spermathecae of all seven *Forcipomyia* females collected from *A. rotunda* flowers and dissected (Figure S3).

### Factors of variation in pollinator communities

The composition of pollinator communities varied significantly among *Aristolochia* species (Deviance = 281, df = 2, p < 0.001; multivariate GLMs implemented with *mvabund*), collection site (Deviance = 514, df = 33, p < 0.001) and month of collection (Deviance = 254, df = 9, p < 0.001). Pairwise comparisons showed significant differences in community composition within all three *Aristolochia* species pairs (adjusted p = 0.001 for all comparisons). The species effect was associated with significant variation in eight pollinator species (including *F. aristolochiae* (p = 0.001), *F. monilicornis* (p = 0.001), *D. punctiventris* (p = 0.003) and *D. turficola* (p = 0.001)), whereas site and month effects were associated with variation in two (*F. aristolochiae* (p = 0.001) and *F. psilonota* (p = 0.003)) and four pollinator species (*F. aristolochiae* (p = 0.001), *F. monilicornis* (p = 0.012), *F. psilonota* (p = 0.002) and *T. ruficeps* (p = 0.001), respectively (univariate tests, adjusted p-values; Table S1). Spatial and temporal variations of pollinator communities of each *Aristolochia* species are represented in Figure S4.

A significant correlation was found between pollinator communities and the geographical distance among sampling sites for the three *Aristolochia* species: *A. rotunda* (Procrustes analysis; r = 0.36, p = 0.001), *A. pistolochia* (r = 0.33, p = 0.001) and *A. clematitis* (r = 0.23, p = 0.002). Pollinator communities changed between the beginning (March) and the end (June) of the main flowering season (Figure S4.b). Ceratopogonidae of the genus *Forcipomyia* were mostly found between March and May, and also between September and October, while Ceratopogonidae of the genus *Dasyhelea* were found in the flowers of *A. clematitis* during all the flowering season. Species belonging to the Chloropidae, and particularly *T. ruficeps*, were mostly found in May and June, at the end of the flowering period of the three *Aristolochia* species.

### Comparisons of floral visitors in synchronopatry

The abundance of some pollinator species in flowers differed significantly among synchronopatric *Aristolochia* species (Figure 1). In synchronopatry between *A. pistolochia* and *A. rotunda, F. aristolochiae* was significantly more abundant in *A. rotunda* flowers than in those of *A. pistolochia* (χ^2^ = 138.7, P < 0.001). In the same situation, *F. monilicornis* was found only in the flowers of *A. pistolochia* (χ^2^ = 8.5, P = 0.004). The abundance of *T. ruficeps* was not significantly different between these two *Aristolochia* species (χ^2^ = 2.1, P = 0.14). In synchronopatry between *A. rotunda* and *A. clematitis*, *F. aristolochiae* was significantly more abundant in the flowers of *A. rotunda* than in those of *A. clematitis* (χ^2^ = 15.3, P < 0.001). *Dasyhelea turficola* (χ^2^ = 8.8, P = 0.003) *and D. punctiventris* (χ^2^ = 2.9, P = 0.09) were found only in the flowers of *A. clematitis*. The number of flowers occupied by *T. ruficeps* (χ^2^ = 33.3, P < 0.001) and by other Chloropidae species considered together (χ^2^ = 24.5, P < 0.001) were also significantly higher in *A. rotunda* than in *A. clematitis*.

**Figure 1:**
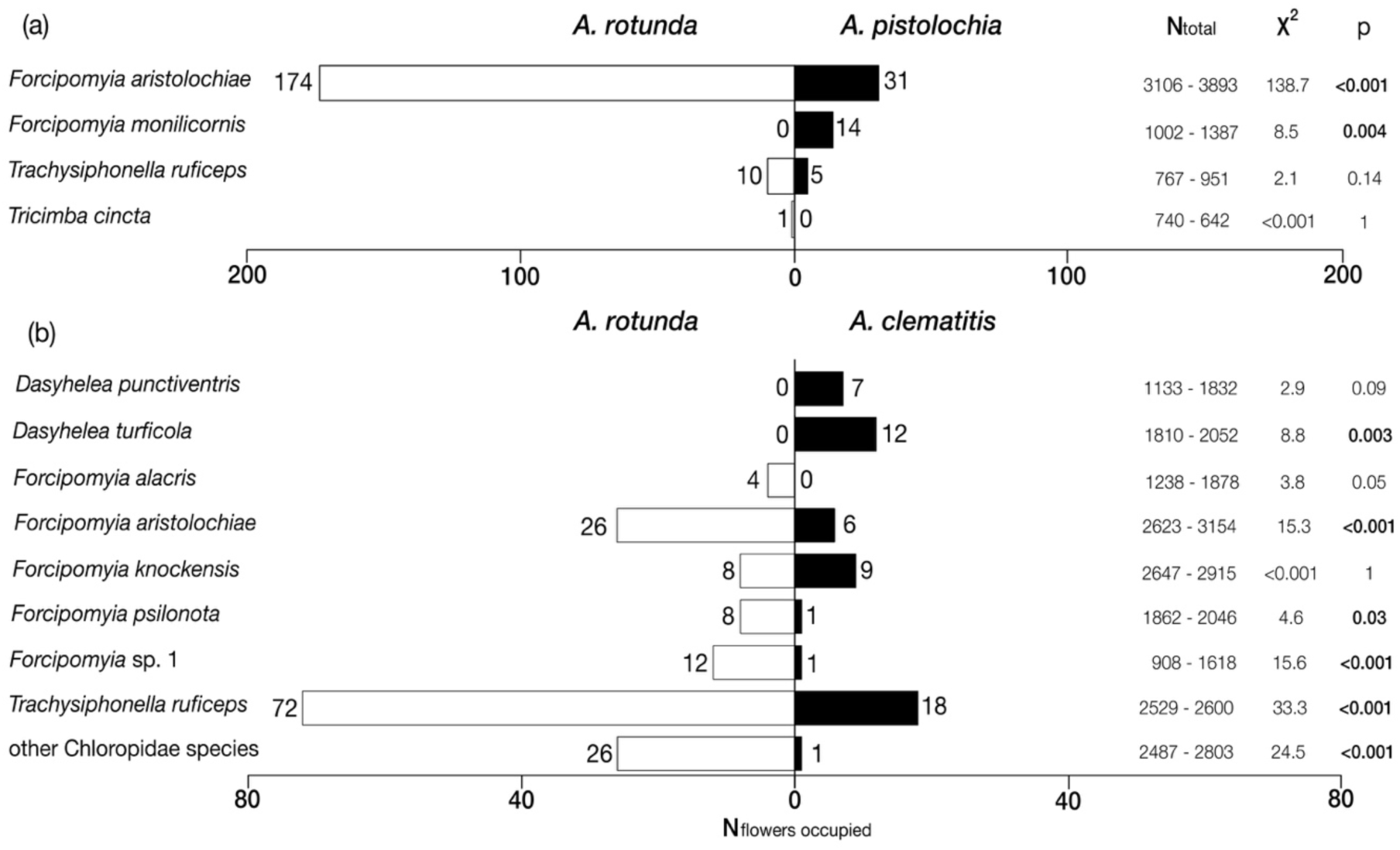
Distribution of the number of flowers occupied by each pollinator species between species of *Aristolochia* in cases of synchronopatry between *A. rotunda* and *A. pistolochia* (a) and between *A. rotunda* and *A. clematitis* (b). N_total_: total number of flowers inspected over multiple sampling sessions, X^2^: Chi-square test value and p-value associated (p), N_flowers occupied_: number of flowers occupied by at least one individual of a pollinator species (see text for details). P-value in bold when < 0.05.

### Floral VOCs

The composition of the extracted floral odour *of A. rotunda* was significantly different from that of *A. pistolochia* (Figure 2, PPLS-DA, Classification error rate = 0%, p = 0.001). Overall, the two *Aristolochia* species differed markedly in the composition of their floral odour, with distinct alkanes, alkenes and aliphatic esters among their most abundant compounds (Table 2; see Table S2 for the complete list of compounds). In detail, the floral odour of *A. rotunda* was dominated by alkanes, which accounted for on average 85% of the blend. The most abundant alkanes included nonane (17.83 ± 10.23%; mean ± SD), decane (30.12 ± 23.67%), dodecane (10.51 ± 4.41%) and tridecane (20.62 ± 13.07%). Esters accounted for 10% of the blend, with compounds such as (*x*)-hex-2-enyl butanoate (7.07 ± 16.99%) among the most abundant, while alkenes accounted for 1.6%, including (*Z*)-tridec-4-ene (0.37 ± 0.24%), (*Z*)-pentadec-6-ene (0.19 ± 0.25%) and (*Z*)-heptadec-8-ene (0.13 ± 0.30%). In contrast, the floral odour of *A. pistolochia* was dominated by alkenes and esters, each of which accounted for more than 30% of the floral blend. Among the most abundant compounds were (*x*)-hex-3-enyl acetate (25.53 ± 19.86%), (*Z*)-pentadec-7-ene (8.91 ± 21.49%), and (*Z*)-pentadec-5-ene (17.01 ± 20.77%). Floral odour of *A. pistolochia* also contained terpenes, which accounted for 24% of the blend, with limonene being the most abundant (11.94 ± 15.46%).

**Figure 2:**
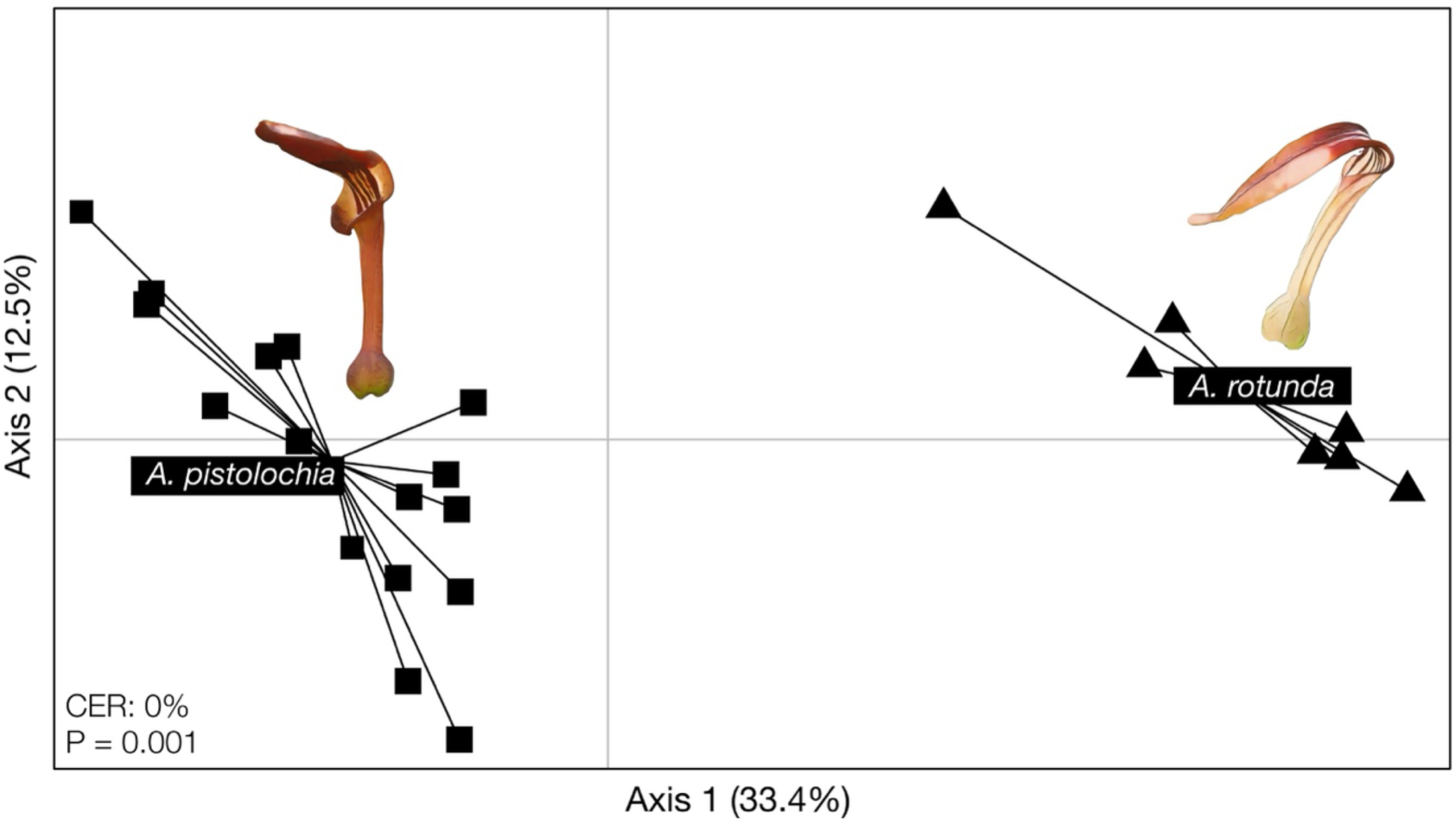
PLS-DA illustrating the differences in the relative proportion of the floral VOCs between seven individuals of *Aristolochia rotunda* and sixteen individuals of *A. pistolochia* in the two first axes. The Classification Error Rate (CER) was 0%, and the p-value was 0.001.

**Table 2:**
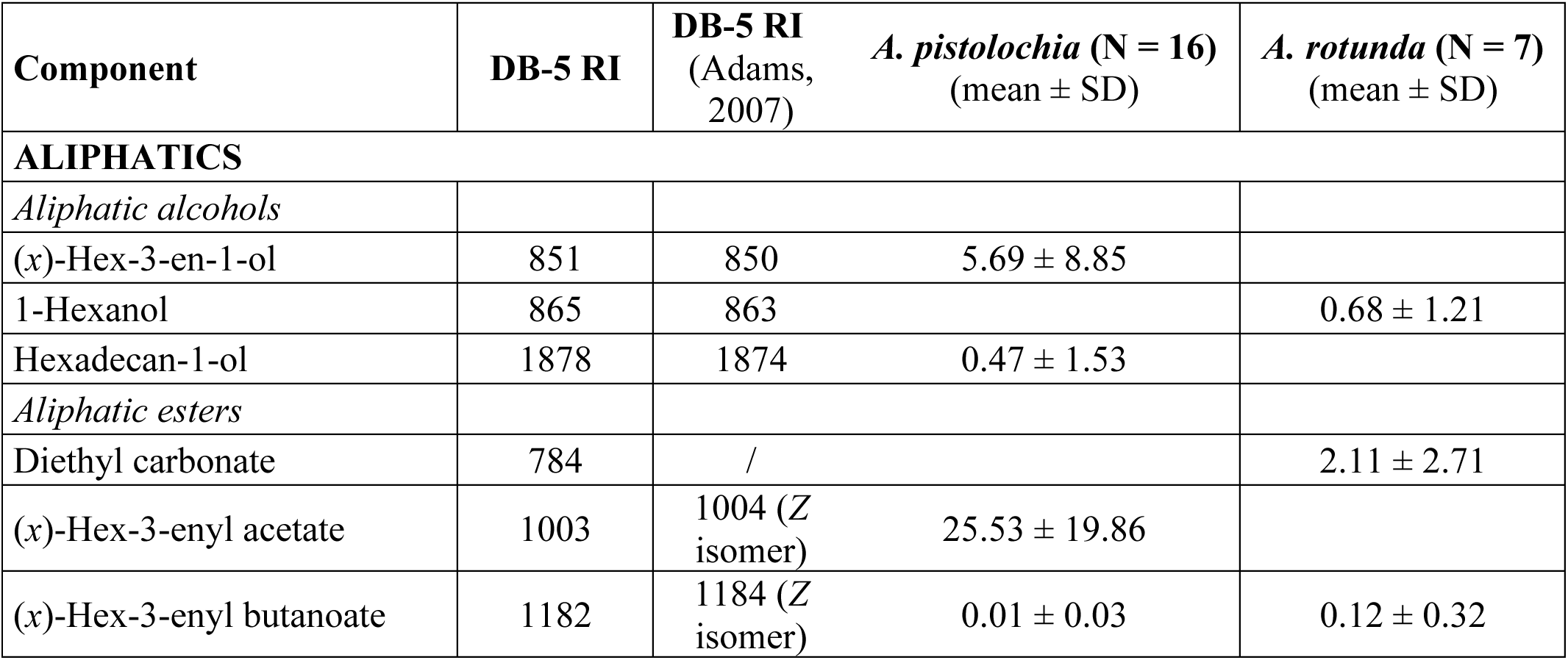
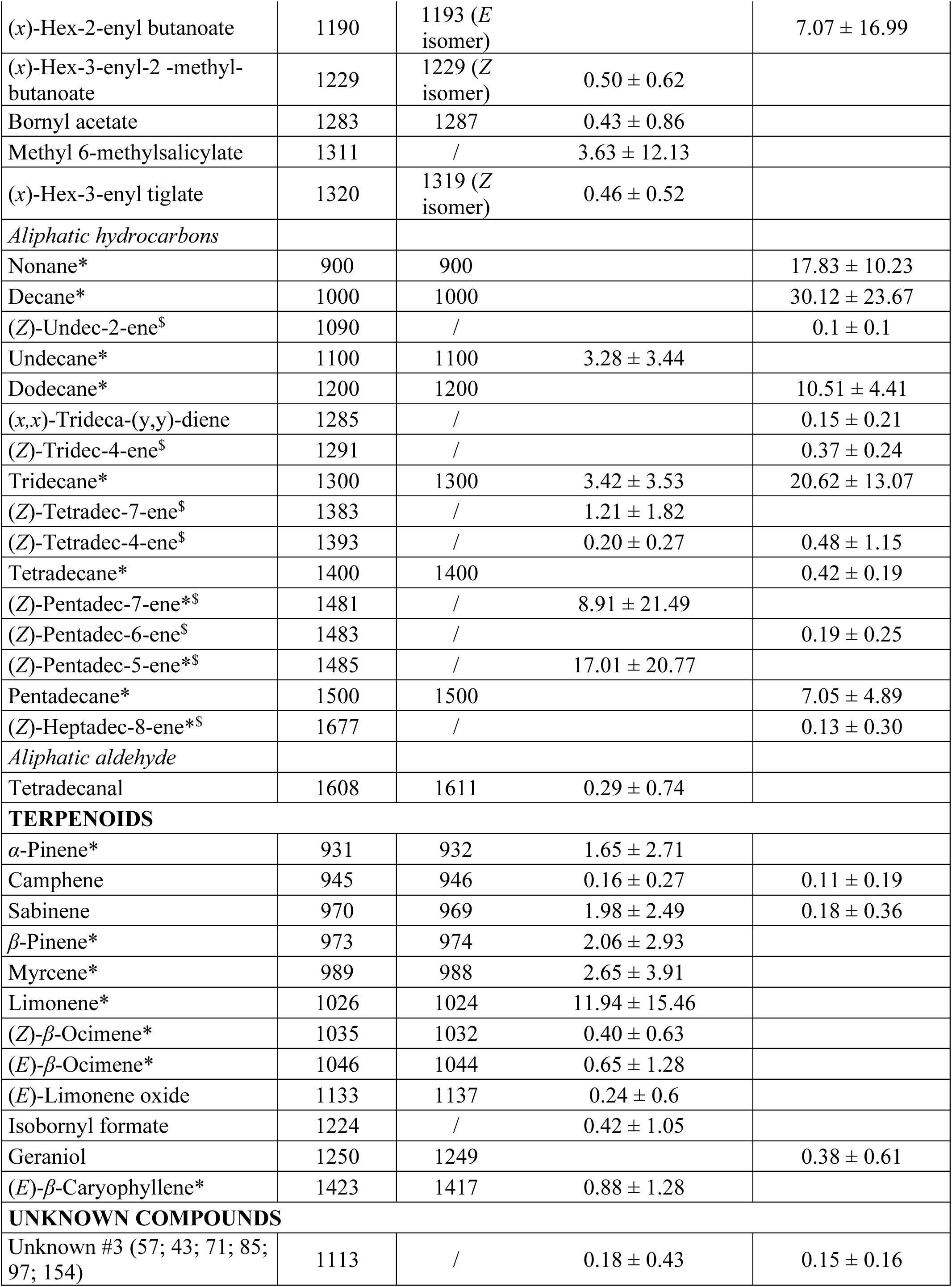
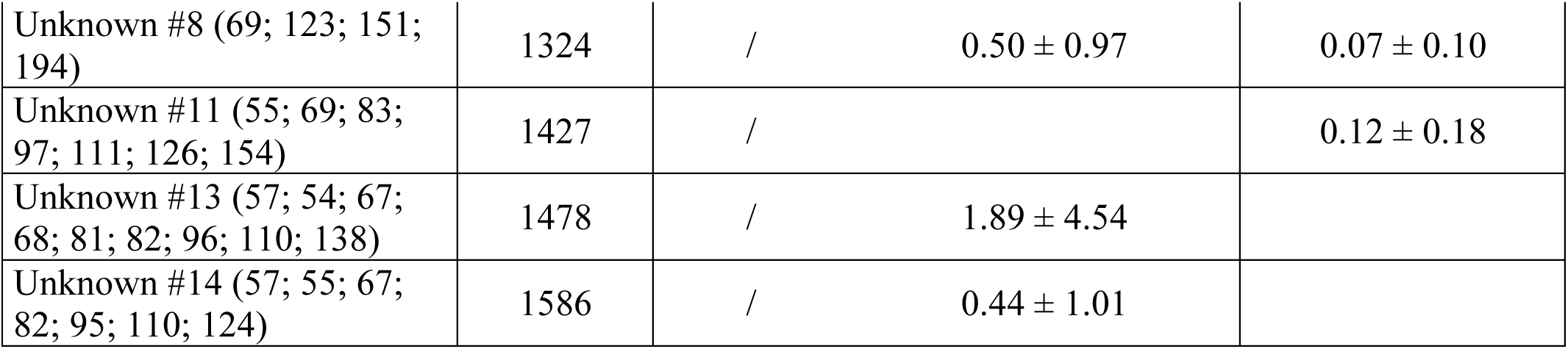
List of the 45 VOCs with the highest proportions among the 89 VOCs detected in the floral odours of *Aristolochia pistolochia* and *A. rotunda*, with their retention indices and mean ± SD relative abundance values (%). * compounds annotations confirmed by comparison with synthetic compounds. ^$^ compounds double bond position and configuration determined by DMDS derivatisation. For the retention index from Adams (2007), when the isomer was uncertain, the closest match was used.

Several compounds that elicited electroantennographic responses in *F. aristolochiae* and *F. monilicornis* (see next section) were detected in fewer than half of the sampled individuals within a given *Aristolochia* species, although they were sometimes present in relatively large amounts. As they did not meet the criterion of being present in more than 50% of the samples, these compounds were excluded from data treatment and subsequent analyses. They included hexyl butanoate, hexyl hexanoate, and one unknown compound (RI = 1080; largest m/z: 58, 72, 114), all three of which were detected in two out of seven (28.5%) individuals of *A. rotunda*. Seventeen compounds (accounting for less than 0.5% and 2% of the blend of *A. rotunda* and *A. pistolochia*, respectively) remained unidentified.

### Pollinator-antennal olfactory detection

A total of 13 compounds induced consistent depolarisation of the antennae of *F*. *aristolochiae* and *F*. *monilicornis* (Figure 3). When *F*. *monilicornis* were exposed to the floral odour of *A. rotunda* and *A. pistolochia*, the set of detected VOCs differed following the differences in the floral VOCs emitted by the two *Aristolochia* species (Figure 3. (a). (c)). Assays performed on *F. monilicornis* (main pollinator of *A. pistolochia*) and *F. aristolochiae* (main pollinator of *A. rotunda*) on the floral odour of *A. rotunda* revealed that almost the same set of VOCs was detected by the two pollinator species in the odour of *A. rotunda* (Figure 3. (a). (b)). The only difference came from hexyl butanoate (or (*x*)-hex-2-enyl butanoate), which was detected by the antennae of *F. monilicornis* but did not elicit a response in most of the assays with *F. aristolochiae*. Due to coelutions of hexyl butanoate and (*x*)-hex-2-enyl butanoate, and of hexyl hexanoate and (*x*)-hex-2-enyl hexanoate, we were not able to discriminate which ones triggered the responses. Electrophysiologically active compounds in the odour of *A. pistolochia* were: undecane, tridecane, (*Z*)-tetradec-7-ene, (*x*)-dec-3-enyl acetate, (*Z*)-pentadec-7-ene, (*Z*)-pentadec-5-ene and pentadecane. In the odour of *A. rotunda*, the electrophysiologically active compounds were: decane, one unknown compound (largest m/z: 58, 72 and 114), undecane, hexyl butanoate (or (*x*)-hex-2-enyl butanoate) (only for *F. monilicornis*), tridecane, hexyl hexanoate (or (*x*)-hex-2-enyl hexanoate), pentadecane, (*Z,Z*)-heptadeca-(6,9)-diene and (*Z*)-heptadec-8-ene. Unfortunately, we did not succeed in obtaining results with *F. aristolochiae* exposed to the odour of *A. pistolochia*, due to the limited number of *F. aristolochiae* individuals tested with the odour of *A. pistolochia* and the low proportion of exploitable electroantennographic responses obtained.

**Figure 3:**
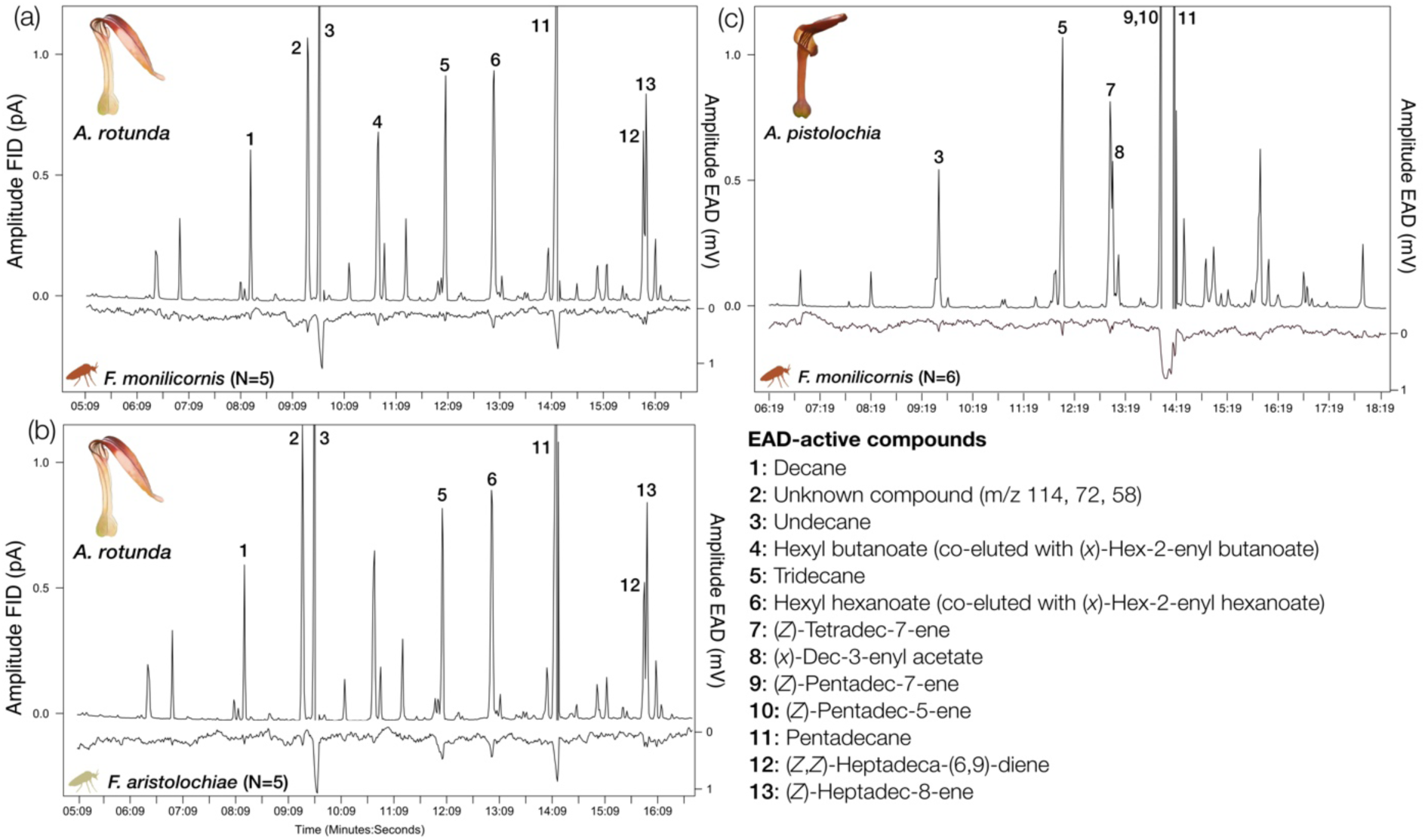
Mean antennal response of (a) *Forcipomyia monilicornis* (main pollinator of *Aristolochia pistolochia*) to the floral odour of *A. rotunda*; (b) *F. aristolochiae* (main pollinator of *A. rotunda*) to the floral odour of *A. rotunda*; (c) *F. monilicornis* (main pollinator of *A. pistolochia*) to the floral odour of *A. pistolochia*. All individuals tested were females. Two compounds are included for the same peak when it was not possible to distinguish between the two due to coelution.

### Field bioassays

Using McPhail traps with hexyl butanoate, 61 individuals of Ceratopogonidae and 627 of Chloropidae were captured, accounting together for almost all the insects captured (> 98.6%). The Ceratopogonidae were all females belonging to *F. monilicornis*. They were captured in only eight traps out of the 127 (6.3%) set, while Chloropidae were found in 77 traps out of the 127 (60.6%) set. Only seven Chloropidae and no Ceratopogonidae were captured in the 60 control traps without odour attractant, while no specimens were captured in the 13 control traps containing only the solvent (acetone).

## Discussion

In the present study, we report that *A. rotunda*, *A. pistolochia* and *A. clematitis* are pollinated by Ceratopogonidae (genera *Forcipomyia* and *Dasyhelea)* and Chloropidae (genera *Trachysiphonella*, *Tricimba*, *Hapleginella*, *Polyodaspis* and *Conioscinella*), and provide a detailed characterisation of these pollinators, including species identity, female reproductive status (i.e., whether females were gravid or not), and pollen loads on both pollinators and stigmas.

Although individuals were found sporadically visiting the other investigated *Aristolochia* species, the main pollinators of *A. pistolochia* (*F. monilicornis*) and *A. clematitis* (*D. turficola* and *D. punctiventris*) were specific pollinators of these two species, according to our data. The main pollinators of *A. rotunda* (*F. aristolochiae* and *T. ruficeps*) were not specific pollinators of this species, although they were preferentially found in *A. rotunda* flowers compared to *A. pistolochia* and *A. clematitis*. Six species of Chloropidae were specific pollinators of *A. rotunda.*, even if they were less abundant than other pollinators of this species. Pollinator communities of the three *Aristolochia* species differed throughout the flowering season and according to geographical location. However, selective or specific visitation of pollinators between pairs of synchronopatric *Aristolochia* species confirmed that traits other than phenology or habitat mediate pollinator specificity. Hence, although species-selective visitation and species-specific pollination occur among the three *Aristolochia* species, and it is likely to be mediated by differences in the floral traits of these plants.

The absence of fruit set through pollinator exclusion in *A. rotunda* and *A. clematitis* indicates that attraction of pollinators is needed to set fruit in these species. In *A. pistolochia*, although one fruit (out of 15 flowers) was produced under the pollinator exclusion treatment, higher fruit set with manual pollination suggests that pollinator attraction also affects reproduction in this species. Our experimental design does not allow determining whether reproductive interference has a cost for reproduction and therefore, whether selection favours the specific attraction of pollinators to achieve or increase fruit set. However, the cost of reproductive interference could be high, as thirteen out of thirty-five (38%) of the collection sites studied have synchronopatric *Aristolochia* species. Thus, relying on different species of pollinators could avoid the cost of reproductive interference.

Interestingly, among the pollinating species, a higher proportion of specimens were detected as pollinators for *A. rotunda* than for the two other *Aristolochia* species. The particularly high proportion of specimens of *F. aristolochiae*—the main pollinator of *A. rotunda*—detected as pollinators in *A. rotunda* may explain this result. Morphological fitting between flies and flowers has been shown to determine the pollinator status of flies in *Aristolochia* (e.g., Brandalise et al., 2026). However, our results on the amount of pollen transported by each pollinator species (see Table 1) suggest that fine-tuned morphological matching may also influence pollen transfer efficiency among pollinators. A comparative study between *A. rotunda* and *A. pistolochia* may be particularly suited to investigate this topic because their floral structure and their respective main pollinators look very similar but, nevertheless, the two plant-pollinator pairs show contrasted efficiency.

In the Mediterranean area, pollination of *Aristolochia* by Ceratopogonidae has only been mentioned for *A. rotunda* and considered secondary, and the genera were not identified (Oelschlagel et al. 2015, 2016). We showed that two additional *Aristolochia* species were pollinated by Ceratopogonidae, and that two genera, *Forcipomyia* and *Dasyhelea*, were involved. At a broader geographic scale, pollination by *Forcipomyia* was previously reported in *Aristolochia bracteolata* Lamk. (Razzak et al., 1992) and in *A. praevenosa* F. Muell. (Sands & Scott, 1998), both located in tropical regions, whereas species of *Dasyhelea* were described as pollinators in *A. contorta* Bunge in Korea (Bang et al., 2026). Although the pollinators were not identified to the genus level, Ceratopogonidae were also described as pollinators in *A. watsonii* Wooton & Standl. in North America (Crosswhite & Crosswhite, 1984). At a broader plant taxonomic scale, Ceratopogonidae were identified as primary pollinators in more than 30 plant species belonging to seven families (Bang et al., 2026; Mochizuki et al., 2025; de Sande et al., 2026; see the review in Vrecko et al., 2025). The Ceratopogonidae pollinators were exclusively, or predominantly, females (de Sande et al., 2026; Vrecko et al., 2025), when sex was recorded. Our results confirm this trend (as for *A. rotunda* and *A. pistolochia*) but also suggest that pollination by Ceratopogonidae may involve both sexes (as for *A. clematitis*). This may reveal distinct pollination strategies involving Ceratopogonidae. Detailed identification of pollinator species and their reproductive status in more plant species is needed to more accurately assess the extent and diversity of pollination strategies involving Ceratopogonidae.

The floral odours of *A. rotunda* and *A. pistolochia* were both dominated by alkanes, alkenes, and aliphatic esters. Many of these compounds elicited electrophysiological responses in the antennae of female *Forcipomyia* pollinators (present study) and some attracted females of a Chloropidae pollinator (Oelschlägel et al. 2015). Among the other *Aristolochia* species for which floral VOCs have been characterised (Alpuente et al., 2023; Johnson & Jürgens, 2010; Martin et al., 2017; Qin et al., 2021; Rupp et al., 2021, 2024; Stashenko et al., 2009), only the floral bouquet of *A. bianorii* was dominated by alkanes, alkenes, and esters (Alpuente et al., 2023). Interestingly, similarly to *A. rotunda*, *A. bianorii* was pollinated by females of Chloropidae and was hypothesised to rely on a kleptomyiophilous strategy for pollination (Alpuente et al., 2023). The presence of these alkanes, alkenes and esters may, then, be related to the attraction of kleptoparasitic flies (Chloropidae) and Ceratopogonidae. Among the other *Aristolochia* species investigated in the literature, which are pollinated by other families of Diptera, alkanes and alkenes have only been detected at a low amounts in the floral odour of *A. gigantea* (Martin et al., 2017). This suggests substantial variability in the composition of floral odours among *Aristolochia* species. As the most abundant compounds have been shown or hypothesised to be associated with pollinator attraction in several *Aristolochia* species (Alpuente et al., 2023; Johnson & Jürgens, 2010; Rupp et al., 2021, 2024), this variability may reflect differences in pollinator-mediated selection among species. Such interspecific variation in floral scent composition, potentially associated with differences in pollination strategies and pollinator taxa, raises questions about the relative contributions of selection and phylogenetic constraints on the evolution of floral specialisation and pollinator specificity.

We highlight significant differences in the composition of the floral odours between *A. rotunda* and *A. pistolochia.* The different electroantennographic responses recorded in *F. monilicornis* exposed to the floral odour of *A. pistolochia* or that of *A. rotunda* indicated that the floral odours of these two species are detected as different floral cues. This suggests that differences in the composition of floral odours could mediate selective pollinator attraction between *A. rotunda* and *A. pistolochia*. Divergence in floral odours is an important factor in mediating reproductive isolation in plant-pollinator interactions (Raguso, 2008; Weber et al., 2018) and is known to play a role in establishing specificity in fly-pollination systems (Kidyoo et al., 2022). However, comparing floral odour composition among plant species is insufficient to confirm that differences in floral bouquets mediate preferential attraction of distinct pollinator species. In the future, this hypothesis will need to be verified experimentally by testing the preferential attraction of distinct pollinator species using floral odour extracts of *A. rotunda* and *A. pistolochia*, or even a mixture of the electrophysiologically active VOCs.

One different electroantennographic response (hexyl butanoate or (*x*)-hex-2-enyl butanoate) was found in the GC-EAD recordings of females of the two pollinator species (*F. aristolochiae* and *F. monilicornis*) when exposed to the odour of *A. rotunda*. This shows that fine differences in olfactory detection capabilities occur between the females of the species of *Forcipomyia*. In the future, experiments with synthetic compounds will be necessary to determine which of the co-eluting esters are electrophysiologically active.

Currently, little is known about how floral VOCs drive the attraction of Ceratopogonidae. Unidentified individuals of the genus *Forcipomyia* were experimentally attracted in the field using a mixture of alkanes (nonane and undecane) and esters (octyl acetate, decyl acetate and methyl-6-methyl salicylate) that reproduced the VOC profile emitted by *Vincetoxicum nakaianum* K. Mochizuki & Ohi-Toma (Apocynaceae), whose flowers are pollinated by kleptoparasitic flies (Chloropidae) (Mochizuki, 2025). The occasional trapping of females of *F. monilicornis* in McPhail traps filled with hexyl butanoate and their absence in control traps (current study) demonstrate that olfactory signals play a role in attraction of this species. Interestingly, hexyl butanoate is among the few VOCs for which the two species differed in detection capability, with *F. aristolochiae* showing neither an antennal nor a behavioural response to this compound, while *F. monilicornis* showed an antennal response and, as confirmed by their response to McPhail traps, also at least a sporadic behavioural response. Specific attraction of females of two *Forcipomyia* species has been shown using traps baited with different aliphatic esters (Sugawara & Muto, 1974). The ability to detect these esters and the triggering of a behavioural response could be very variable among all the *Forcipomyia* species. The alkanes and alkenes detected by *F. aristolochiae* and *F. monilicornis* in the floral odours of *A. rotunda* and *A. pistolochia* are also found in the floral odour of *Theobroma cacao* L. (Malvaceae) (Arnold et al., 2019). Further experimentation is required to better understand how floral VOCs drive the specific behavioural attraction of the pollinating ceratopogonid flies, particularly in comparison to other potential attractive signals, such as visual cues.

Interestingly, *T. ruficeps* (Chloropidae), which was previously identified as the main pollinator of *A. rotunda* in Croatia (Oelschlägel et al., 2015, 2016), plays a secondary role in pollination of this species at our study sites. This difference may reflect geographical variation in the composition of pollinator assemblages. Alternatively, since the pollinator sampling in Croatia was conducted in May only (Oelschlägel et al., 2015, 2016) the sampling time may have been outside the period when pollinating ceratopogonids are most active (Figure S3). The high variability in abundance of hexyl butanoate, (*x*)-hex-2-enyl butanoate, hexyl hexanoate and (*x*)-hex-2-enyl hexanoate among individual flowers found in our results was also highlighted by Oelschlägel et al. (2015). Although these compounds were electrophysiologically detected by *T. ruficeps* (Oelschlägel et al., 2015), *F. aristolochiae* (the main pollinator of *A. rotunda* in our study) does not appear to detect hexyl butanoate or (*x*)-hex-2-enyl butanoate. Different VOCs can act as cues advertising different mimicked resources for different pollinator taxa. Thus, the mimicry of injured true bugs, as described by Oelschlägel et al. (2015), should only be directly attributed to the relation between *A. rotunda* and *T. ruficeps* and, without further evidence, should not be extrapolated to the plant’s pollination strategy with regard to other pollinators, such as Ceratopogonidae.

The differences in identity, sex and reproductive status of pollinators among the three *Aristolochia* species could be explained by differences in the pollination strategies employed by these plants. In *A. pistolochia* and *A. rotunda*, almost all the individuals of Ceratopogonidae pollinators were females, and neither eggs nor gravid individuals were observed. In addition, sperm was found in the spermathecae of all seven *Forcipomyia* females collected from *A. rotunda* flowers. As the females had already mated but were not gravid, the resource sought in the flowers was likely neither a brood site nor a sexual partner. As females of many species of *Forcipomyia* need to feed on insect haemolymph to mature their eggs (Kirk-Spriggs & Sinclair, 2017), our hypothesis regarding pollination strategy was that the two *Aristolochia* species mimic the insect host of their respective *Forcipomyia* pollinators (following the rationale in Vrecko et al., 2025). This hypothesis was reinforced by the previous finding that flowers of *A. rotunda* attract *T. ruficeps* by mimicking the odour of injured bugs (Oelschlägel et al., 2015). In addition, other plant species belonging to a different plant family attracting kleptoparasitic flies by mimicking insects were also shown to attract Ceratopogonidae (Heiduk et al., 2023; Mochizuki, 2025). The pollinator-detected alkenes and esters in the odour of *A. rotunda* and *A. pistolochia* are known to function as communication or defence compounds in various insect taxa. (*Z*)-Tridec-5-ene, (*Z,Z*)-heptadeca-(6,9)-diene and (*Z*)-heptadec-8-ene have been identified as defensive compounds in Carabidae (Coleoptera) (Moore, 1979), whereas hexyl hexanoate, hexyl butanoate and (*x*)-hex-2-enyl butanoate are known as defensive compounds in some families of bugs (Heteroptera) and in Staphylinidae (Coleoptera) (Yang et al., 2014, 2015). In the floral odour of *A. pistolochia*, (*Z*)-pentadec-7-ene and (*Z*)-pentadec-5-ene have also detected in the Dufour’s gland of some species of Formicidae (Hymenoptera) (Bagnères et al., 1991). It is unknown whether all these compounds are detected by males, or whether the olfactory-active VOCs that were characterised are specifically detected by females to seek their host (or any resource on which they collect the nutrients they need for egg maturation). It is also possible that these compounds do not mimic a specific host insect, but rather that these *Aristolochia* species exploit a perceptual bias in the Ceratopogonidae associated with the presence of a reward, as has been suggested in other types of plant-insect interactions (Ruxton & Schaefer, 2011). However, the *Forcipomyia* pollinators of the two *Aristolochia* species belong to the subgenera *Euprojoannisia* (*F*. *alacris*), *Panhelea* (*F*. *aristolochiae*), *Synthyridomyia* (*F. knockensis*) and *Thyridomyia* (*F. monilicornis*), which have never been observed to suck blood or haemolymph but are known to be important flower visitors (de Meillon & Wirth, 1991), calling into question the validity of the hypothesis of host mimicry. Further evaluation of this hypothesis depends on understanding the life cycle of *Forcipomyia* pollinators.

Finally, the specific pollinators of *A. clematitis* were males and females of two species of *Dasyhelea* (Ceratopogonidae): *D. turficola* and *D. punctiventris.* A balanced sex-ratio was observed in the individuals of *Dasyhelea* collected in the flowers of *A. clematitis*, and some females possibly had maturing eggs. The attraction of males and possibly gravid females makes the hypothesis of exploitation by the plant species of the nutrient-seeking behaviour needed for egg maturation very unlikely. Furthermore, unlike *Forcipomyia*, adult *Dasyhelea* females have reduced mouthparts and do not bite vertebrates or invertebrates (Kirk-Spriggs & Sinclair, 2017). The deceptive or mutualistic nature of pollination in *A. clematitis* remains to be elucidated.

## Acknowledgments

We thank the PACE (*Plateforme d’Analyses Chimiques en Ecologie*) of the “Montpellier Ecologie Environnement Biodiversité” (MEEB) laboratory, UAR 2040 CNRS-UM, hosted by the CEFE, for providing access to its facilities and support for the chemical analyses. We are grateful to Marie-Pierre Dubois for help during the experiments conducted at the GEMEX (*Génomique, Ecologie Moléculaire, Evolution Expérimentale*) technical facility, and Pauline Durbin and Thierry Mathieu to the *Plateforme des terrains d’expériences* of the *Centre d’Ecologie Fonctionnelle et Evolutive*, with the support of LabEx CeMEB, an ANR *Investissements d’avenir* program (ANR-10-LABX-04-01). We are very thankful to Stefan Schulz (Technische Universität Braunschweig) for giving us the alkadiene that we needed for our experiments and to Mathilde Dufaÿ, Pierre-Olivier Cheptou and Finn Kjellberg, all at the CEFE, for the help provided in the conceptual design of the study. We thank the *Conservatoire du Littoral* and the *Conservatoire d’Espaces Naturels* and the people working in these agencies for providing access to study sites. We thank Christophe Bernier for providing information on the location of *Aristolochia* in the studied area and for providing access to study sites.

## Data, scripts, code, and supplementary information availability

Datasets, code and supporting material (Supporting Method and Figures S1-S4) are available at the following DOI: https://doi.org/10.48579/PRO/ZQQSFW

